# Bone marrow mesenchymal stem cells from osteoporotic patients do not show altered mitochondrial energetics or ultrastructure

**DOI:** 10.1101/046300

**Authors:** Clemens W.I. Moll, Richard A. Lindtner, Georg F. Vogel, Martina Brunner, Karin Gutleben, Michael Blauth, Gerhard Krumschnabel, Hannes L. Ebner

## Abstract

As a consequence of the ongoing demographic change, osteoporosis is considered as one of the mayor challenges for the health care system of the 21^st^ century. However, the exact etiology of osteoporosis is far from being understood. Some evidence suggests that changes in stem cell metabolism might contribute to development of the disease. Therefore we evaluated whether differences of the morphology and/or the energy metabolism of mitochondria can be observed between human bone marrow derived mesenchymal stem cells obtained from osteoporotic patients as compared to non-osteoporotic controls.

Mesenchymal stem cells were isolated from the bone marrow of senile osteoporotic and non osteoporotic patients, osteoporosis being assessed by dual energy X-ray absorptiometry. We then confirmed the stemness of the cells by FACS analysis of the expression of surface markers and by conducting multi-lineage differentiation experiments. And we finally investigated mitochondrial morphology and function with electron microscopy of cryo-fixed samples and by high-resolution respirometry, respectively. In addition we compared the energy metabolism of the stem cells to those of the osteosarcoma cell line MG-63.

The data show, for the first time, the applicability on stem cells of the methods used here. Furthermore, our results indicated that there are no obvious differences detectable in mitochondrial morphology between cells from osteoporotic and non osteoporotic donors and that these cells also seem to be energetically indistinguishable with unchanged rates of routine respiration and respiratory capacity as well as unaltered oxygen consumption rates linked to different respiratory complexes. In summary, we could not detect any evidence indicating major changes of mitochondrial features in cells from osteoporotic patients.

## 1. Introduction

Osteoporosis is characterized as a typically aging related disease associated with low bone mass and a resulting elevated risk for bone fractures. Accordingly, low trauma fractures are often the reason for further evaluation of the bone which may then lead to the diagnosis of osteoporosis. The gold standard for diagnosis and follow-up controls is Dual-energy X-ray absorptiometry (DXA) scanning which measures the bone mineral density [1]. According to the World Health Organization a bone mineral density ≥2.5 SD below the normal mean for young-adult women classifies for established osteoporosis. Diagnostically osteoporosis is divided into primary and secondary osteoporosis, where in the latter form no specific underlying disease as a reason for diminished bone mass is detectable. Primary osteoporosis may be distinguished into juvenile and idiopathic forms, the latter mostly comprising postmenopausal and age-associated osteoporosis with over 30% of all postmenopausal women in Europe and the United States suffering from this chronic and progressive process. With a lifetime risk of sustaining one or more fragility fractures in at least 40%, the socioeconomic burden is huge [2] and osteoporosis therapy reducing fracture risk and preventing further bone mass loss is therefore clearly desirable. However, to develop therapy strategies, a better understanding of the pathophysiology is pivotal. Only in recent years we have started to understand the complexity of this metabolic illness with multiple pathomechanisms. Among the principle issues detected is an apparent imbalance between bone remodeling and bone resorption with an altered function and number of osteoblast and osteoclasts being discussed to be the cause [3]. This results not only in a deterioration of the microarchitecture of affected bone but also in a decrease of overall bone mass. Noteworthy, results of mesenchymal stem cell studies indicate that human bone marrow derived mesenchymal stem cells (hBMSC) may be key players in this regard [4]. These cells are the progenitors, among others, of the cells differentiating to osteoblasts, and they are therefore thought to hold great potential in regenerative medicine, including the potential treatment of osteoporosis [5]. Further, a growing body of evidence has implicated a potential role of alterations in mitochondrial DNA (mtDNA) in the development of osteoporosis [6,7]. Given the importance of mitochondria in cellular energy metabolism and the fact that components of the electron transfer system are encoded by mtDNA, it seems conceivable that altered mitochondrial energy metabolism may be associated with the osteoporotic phenotype [8]. In line, stem cells are believed to undergo dramatic metabolic changes during differentiation, involving changes of mitochondrial arrangement and abundance [9]. Altogether, this led us to speculate that potential alterations in the energy metabolism of mesenchymal stem cells could be an important aspect in this scenario, as this may affect the above noted imbalance between bone-forming and removing cells both by altering cell numbers and activity. Therefore we investigated the most crucial parameters reflecting mitochondrial function, oxygen consumption and mitochondrial morphology, using, in the first study of this kind, hBMSC from non-osteoporotic and osteoporotic donors and comparing their respirometric characteristics with those of a commercially available osteosarcoma cell line (MG-63 cells).

## 2. Material und Methods

### 2.1 Patients and ethical approval

The study was approved by the ethics committee of the medical university of Innsbruck. Written informed consent was obtained from all the participants. hBMSC were isolated from 13 aged donors undergoing routine surgical procedures for treatment of proximal femur fracture at the University Hospital Innsbruck, Department of Trauma Surgery, Austria (Table 1). The inclusion criteria were that patients were older than 65 years and were undergoing a standard surgical procedure for hip fracture. Osteoporosis was confirmed in 7 of these patients using dual energy X-ray absorptiometry (DXA; Hologic QDR 4500). The patients showing a T-score of ‐2.5 standard deviations (SD) or less at the contra lateral hip, the femur and/or lumbar spine were considered as osteoporotic, whereas the 6 patients showing a T-score higher than ‐1 and lower than 2.5 were considered as non-osteoporotic.

**Table 1:** Classification of patients with age, sex and Dual-energy X-ray absorptiometry scanning score.

| Patient | age | sex | DXA<sup>1</sup> hip | DXA fem. | DXA spine | OP/nOP |
| --- | --- | --- | --- | --- | --- | --- |
| 1 | 89 | F | -1,8 | -2,5 | -0,8 | OP<sup>3</sup> |
| 2 | 84 | m | 0,7 | 0,1 | 1,6 | nOP<sup>3</sup> |
| 3 | 84 | F | -2,4 | -2,4 | -3 | OP |
| 4 | 71 | m | -0,4 | -0,5 | 0,8 | nOP |
| 5 | 84 | F | prost.<sup>2</sup> | prost. | -5,1 | OP |
| 6 | 74 | m | 0,1 | -0,6 | 0,6 | nOP |
| 7 | 68 | m | 0,3 | -0,2 | -0,1 | nOP |
| 8 | 90 | F | -2,7 | -3 | -1,5 | OP |
| 9 | 69 | m | -0,4 | -1,3 | -0,4 | nOP |
| 10 | 93 | F | -3,5 | -3,6 | -2,3 | OP |
| 11 | 72 | m | -3,9 | -4,3 | -5,6 | OP |
| 12 | 74 | F | -1,9 | -2,5 | -1,7 | OP |
| 13 | 66 | F | -0,4 | -0,2 | -0,9 | nOP |

<sup>1</sup> DXA denotes Dual-energy X-ray absorptiometry scanning by which bone mineral density was assessed; DXA hip, fem., spine denote scores obtained on measurements at the contra lateral hip, the femur and the lumbar spine, respectively.
<sup>2</sup> prost. indicates that a measurement was not possible due to already implanted prosthesis;
<sup>3</sup>OP and nOP signify patients classified as osteoporotic and non-osteoporotic, respectively.

### 2.2. Cell culture

Bone marrow was harvested from the femoral medullary cavity of patients undergoing routine surgery and hBMSC isolated according a well-established protocol [10]. Purified hBMSCs were maintained at 37 °C, in 5% CO_2_ and 98% humidity in normal growth medium consisting of Dulbecco’s modified eagle medium (DMEM-low glucose, with L-Glutamine) (PAA Laboratories GmbH, Pasching, Austria), supplemented with 10% fetal bovine serum (FBS) (Biowest, Nuaillé, France) and 1% penicillin (50 units/ml) and streptomycin (50 μg/ml) (Gibco, Vienna, Austria). The medium was changed twice a week. Cells were used from passage 2 - 4. For respirometric experiments hBMSCs were cultured in 75 cm Cellstar flasks (Greiner Bio-One, Frickenhausen, Germany) up to confluency. Furthermore, the osteosarcoma cell line MG-63 was cultured under the same conditions and used for comparison in respirometry. For electron microscopy hBMSC were cultured on sapphire discs as described by Vogel G.F. et.al [11].

### 2.3. Cellular stemness

Sternness of BMSC was assessed according the minimal criteria for defining multipotent mesenchymal stromal cells defined by the ISCT (International Society for Cellular Therapy)[12]. In short this included FACS analysis of cells stained with phycoerythrin labeled antibodies (Biozym scientific GmbH, Oldendorf, Germany) specific for CD90, CD105, CD166, CD34, CD14, CD45 and HLA-DR. Antibody specificity was confirmed through the use of appropriate phycoerythrin-labeled isotype control antibodies. Flow cytometry was carried out on a FACS Calibur (BD Biosciences, Schwechat, Austria) and staining analyzed using BD CellQuest™ Pro‐ and FlowJo™ software. Furthermore, plastic adherence was shown on the 75 cm^2^ flasks during the first passages and, finally, trilineage differentiation along the adipo‐, chondro‐ and osteogenic lineages was confirmed by using standard differentiation media and procedures well established in our lab [10].

Osteogenic differentiation was induced by incubating hBMSC with osteogenic induction medium (OM) consisting of normal growth medium supplemented with 50 μM L-Ascorbic acid 2-phosphate (Sigma-Aldrich), 10 mM β-glycerophosphate (Sigma-Aldrich) and 100 nM dexamethasone (Sigma-Aldrich). The mineralized matrix being formed upon induction was visualized by Alizarin Red S staining at day 21 of differentiation.

Adipogenic differentiation was induced by incubating hBMSC for 48h in adipogenic induction medium, consisting of Dulbecco’s modified eagle medium (DMEM-high glucose with L-glutamine and with sodium pyruvate), 10% FBS and 1% pen/strep, supplemented with 1 μM dexamethasone, 10 μg/ml insulin, 120 μM indomethacin and 0.5 mM isobutylmethylxanthine (IBMX). Following this 48h induction period the medium was exchanged for adipogenic maintenance medium (induction medium minus IBMX) and cells were cultured for another 14 days. These cells showed intracellular lipid droplets, similar to adipocytes, and these were visualized by Oil red O staining.

Chondrogenic differentiation was carried out by plating hBMSC in 5 μl droplets at a density of 1,6^*^10^7^ ml^-1^ micro mass cultures and differentiating them according the instructions given by the STEMPRO^®^ Chondrogenesis Differentiation Kit (Gibco Life Technologies, Paisley, UK) for up to 21 days. Visualization of newly formed proteoglycans was carried out by staining the pellets with Alcian blue. Pictures were taken on a LEICA M50 stereo microscope or a LEICA DMI300B and images captured with a LEICA 450 DFC camera and processed with Adobe Photoshop CS5 software.

### 2.4. Viability

For the assessment of cell viability cells were cultured on six well plates, de-attached by trypsinization and resuspended in 200 μl PBS in a standard FACS tube. These suspended cells were incubated with 1 μg/ml propidium iodide (AbD Serotec, Puchheim, Germany) for 30 min and propidium iodide exclusion was then measured on a FACS Calibur. Cells were used for experiments when showing a viability ≥90%.

### 2.5. Electron microscopy

TEM (transmission electron microscopy) samples were subjected to rapid cryo-immobilisation by means of high-pressure freezing instead of chemical fixation, which greatly improves the preservation of the mitochondrial ultrastructure. In brief, cells were cultured on sapphire discs and were cryo fixed by high-pressure freezing followed by freeze-substitution and araldite resin embedding for morphological-ultrastructural analysis as described earlier [11]. Ultra-thin sections were analyzed on a Philips CM120 EM (F.E.I., Eindhoven, Netherlands). Images were taken with a MORADA digital camera (SIS, Münster, Germany). ADOBE photoshop software (Version CS6) was used to adjust contrast and brightness of images which were further processed by grey scale modification and high-pass filtering to increase contrast and sharpness.

### 2.6. Electron tomography

For tomography, 250-nm thick-sections were cut and placed on formvar coated slots and coated with 10-nm fiducial gold. TEM tomography was performed on a Tecnai G2 (F.E.I., Eindhoven, Netherlands) microscope at 200KV. Dual-axis (0° and 90°) tilt series images were recorded at 6500x magnification from ‐65° to 65° with 1° increments using Xplore3D automated tomography software (F.E.I., Eindhoven, Netherlands) at a binning of 2 with a 4kx4k pixel camera (Eagle, F.E.I., Netherlands). Reconstruction of back-projected tomography volumes was done using IMOD, contours were manually assigned using 3dmod and movies were made from exported IMOD images using Fiji software (public domain, http://fiji.sc/Fiji).

### 2.7. High-resolution respirometry

Respirometry of intact and permeabilized cells was performed by high-resolution respirometry using the Oroboros Oxygraph-2k with dedicated software DatLab 5 (Oroboros Instruments, Corp., Innsbruck, Austria) following protocols described by Pesta and Gnaiger[13]. Cells were harvested by trypsination, counted and immediately used for respirometry at a temperature of 37°C with media equilibrated to air saturation. For measuring respiration in intact cells, 0.2^*^10^6^ cells ml^-1^ (hBMSC) or 0.3^*^10^6^ cells ml^-1^ (MG-63) were injected into the 2 ml measuring chamber containing cell culture medium and examined applying a coupling control protocol [13]. This protocol is designed to characterize the main functional aspects of cellular respiration in intact cells in a physiologically relevant setting, i.e. under conditions where the cellular environment, by virtue of selecting the standard culture medium, resembles that experienced *in vivo.* Thus, mitochondria do neither experience saturating substrate supply eliciting maximum respiratory rates, nor are they challenged by extreme cellular ATP demand, and overall mitochondria are thus respiring at a rate that can be referred to as ROUTINE respiration. The initial assessment of the ROUTINE rate of respiration was followed by addition of 1 μg^*^ml^-1^ of the ATP synthase inhibitor oligomycin to obtain LEAK respiration. LEAK respiration represents the component of respiration supporting the maintenance of mitochondrial membrane potential by compensating primarily for dissipative proton flux across the inner membrane which is not directly related to ATP production. Next, the uncoupler carbonyl cyanide m-chloro phenyl hydrazone (CCCP) was sequentially titrated until the maximum rate of respiration was observed reflecting the electron transfer system capacity, ETS, under the conditions applied. Finally, 0.5 μM complex-I inhibitor rotenone and 2.5 μM complex-III inhibitor antimycin A were added to obtain a measure of residual oxygen consumption, ROX, not linked to activity of the electron transfer system.

Respiration of permeabilized cells was conducted in medium MiR05 (0.5 mM EGTA, 3 mM MgCl_2_·6H_2_O, 60 mM K-lactobionate, 20 mM taurine, 10 mM KH_2_PO_4_, 20 mM Hepes, 110 mM sucrose, and 1 mg^*^ml^-1^ bovine serum albumin, pH 7.1 at 30°C) after establishing the optimum concentration for digitonin of 7.5 μg^*^ml^-1^ for cell permeabilization in preliminary experiments. In this protocol cells were first permeabilized with digitonin to enable unlimited access of externally added substrates and inhibitors to mitochondria. Thus, activity of individual respiratory complexes may be assessed and their operation under saturating substrate conditions can be evaluated. At first, substrates supporting complex-I linked respiration were added (5 mM pyruvate, 0.5 mM malate, 10 mM glutamate) to observe LEAK respiration. Next, a saturating concentration of 2.5 mM ADP was added to induce primarily complex-I supported OXPHOS capacity, followed by injection of 10 mM succinate resulting in convergent complex-I and ‐II supported OXPHOS capacity. Subsequent titration of CCCP to maximum ETS respiration was followed by addition of rotenone to obtain complex II-linked respiration and antimycin A to determine ROX. Addition of cytochrome c in the OXPHOS state to test for mitochondrial membrane integrity resulted in no significant increase of respiration in these experiments.

## 3. Results

### 3.1. Characterization of hBMSC from senile osteoporotic and non-osteoporotic patients

hBMSC were isolated from 13 senile patients (≥ 66 years) of which 7 fulfilled our criteria for the osteoporosis group with bone mineral density ≥2.5 SD below the normal mean of healthy adult women, whereas 6 were considered as not having any bone defect. Age, sex and T scores derived from DXA measurements are summarized in Table 1. From this overview it is evident that osteoporotic patients were 84 ± 8 years (SD) in average and mainly female (6 out of 7) while non-osteoporotic patients were 72 ± 6 years and mainly male (5 out of 6), in part reflecting, although clearly overstressing, the uneven incidence of osteoporosis in women and men.

The investigated cells isolated from the above described donors showed high expression levels of known surface markers related to mesenchymal stem cells CD166, CD105 and CD90 (Fig. 1A, panels b-d) compared to the unstained negative control (Fig. 1A, panel a), whereas the expression levels for markers of the hematopoietic cell line CD45, CD14 and CD34 as well as for HLA-DR (Fig. 1A, panels e-h) were negligible (negative). Viability of isolated cells assessed with PI staining amounted to 96 ± 3.9% (n = 4).

**Fig. 1:**
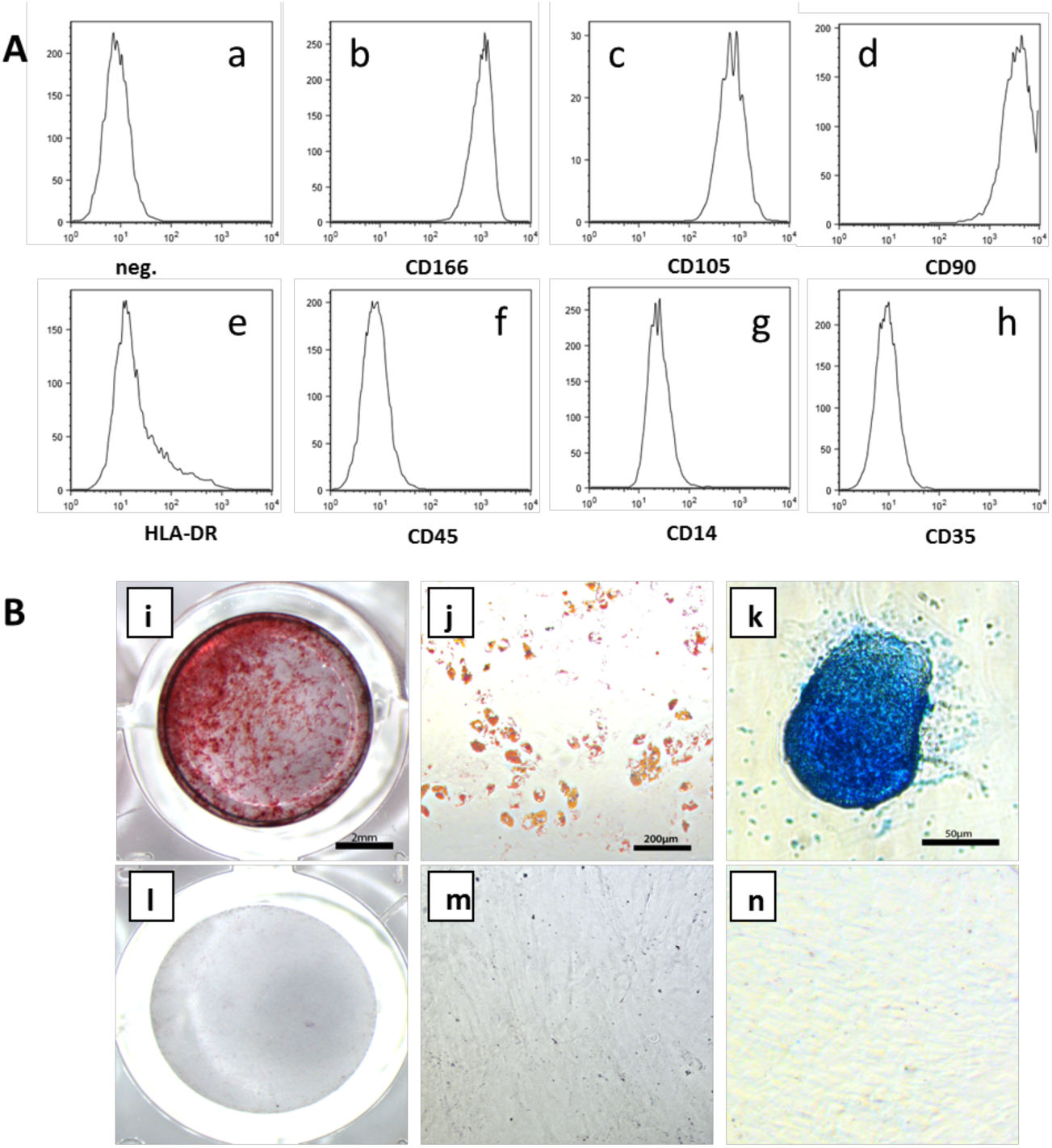
Characterization of hBMSC isolated from aged patients. A: FACS analysis of surface marker expression. Image a, negative control (unstained cells), images b-h, expression pattern of antigens known to be characteristic for mesenchymal stem cells, CD166, CD105 and CD90, and the lack of expression of HLA-DR and the hematopoietic cell line markers CD45, CD14 and CD34. B: Trilineage differentiation of hBMSC. Osteogenic differentiation after 21d in osteogenic induction medium. Staining of mineralized matrix was performed with Alizarin Red S (i, l). Newly formed lipid droplets shown after 16d of adipogenic induction and visualized by Oil Red O staining (j, m). Chondrogenic differentiation shown on a pellet culture incubated for 21d in chondrogenic induction medium and stained for proteoglycans with Alcian blue (k, n). The upper panel shows cells stained after incubation in differentiation medium, the lower panel displays cells incubated in normal growth medium.

hBMSC showed the expected differentiation pattern along all three pathways of the osteogenic-, the adipogenic-, and the chondrogenic lineage. Alizarin Red S staining was performed to show the Ca^2+^ and hydroxyapatite rich matrix conventionally formed by osteoblasts (Fig. 1B, panel i). In contrast, upon induction of adipogenesis the characteristic formation of lipid droplets appeared in bright red colour after staining the differentiating hBMSC with Oil Red O at day 14 of induction (Fig. 1B, panel j). Pellet cultures differentiated for 21 days along the chondrogenic lineage showed a positive staining for proteoglycans following Alcian blue staining from chondrocyte like cells (Fig. 1B, panel k). Negative controls performed on the same well-plate never stained positive for any dye following the same incubation periods and incubation in normal grow medium followed by the same staining procedures. (Fig. 1B, panels l, m, n).

### 3.2. Morphological characterization of hBMSC

Transmission electron microscopic images were used to assess potential changes in mitochondrial morphology and/or distribution. These parameters were not strictly quantified but were independently assessed qualitatively by two researchers experienced with electron microscopic analyses. This analysis of high pressure frozen, rehydrated samples showed a predominantly perinuclear localization of the mitochondria in cells from osteoporotic (OP) and non osteoporotic (nOP) donors as described earlier by Lonergan et. al. for various stem cell lines [9]. The number of mitochondria visible in the cultured cells appeared unaltered in OP cells (Fig 2A, D) and is thus indicative of a normal metabolic activity of the cells. Further, the organelles were observed to be organized in an extended tubular, threadlike meshwork with no signs of altered morphology such as mitochondrial swelling and altered number or size of cristae, indicating appropriate functionality of the investigated organelles (Fig 2C, F). Interestingly, in cells from both, OP and nOP donors, a high number of lyso-autophagosomes could be observed, which is usually considered a sign of unhealthy cells and high membrane turnover. In line, the abundantly occurring rough endoplasmic reticulum underpins this assumption. Overall, electron microscopic analysis did not indicate any apparent differences in the ultrastructure between cells from OP and nOP donors.

**Fig. 2:**
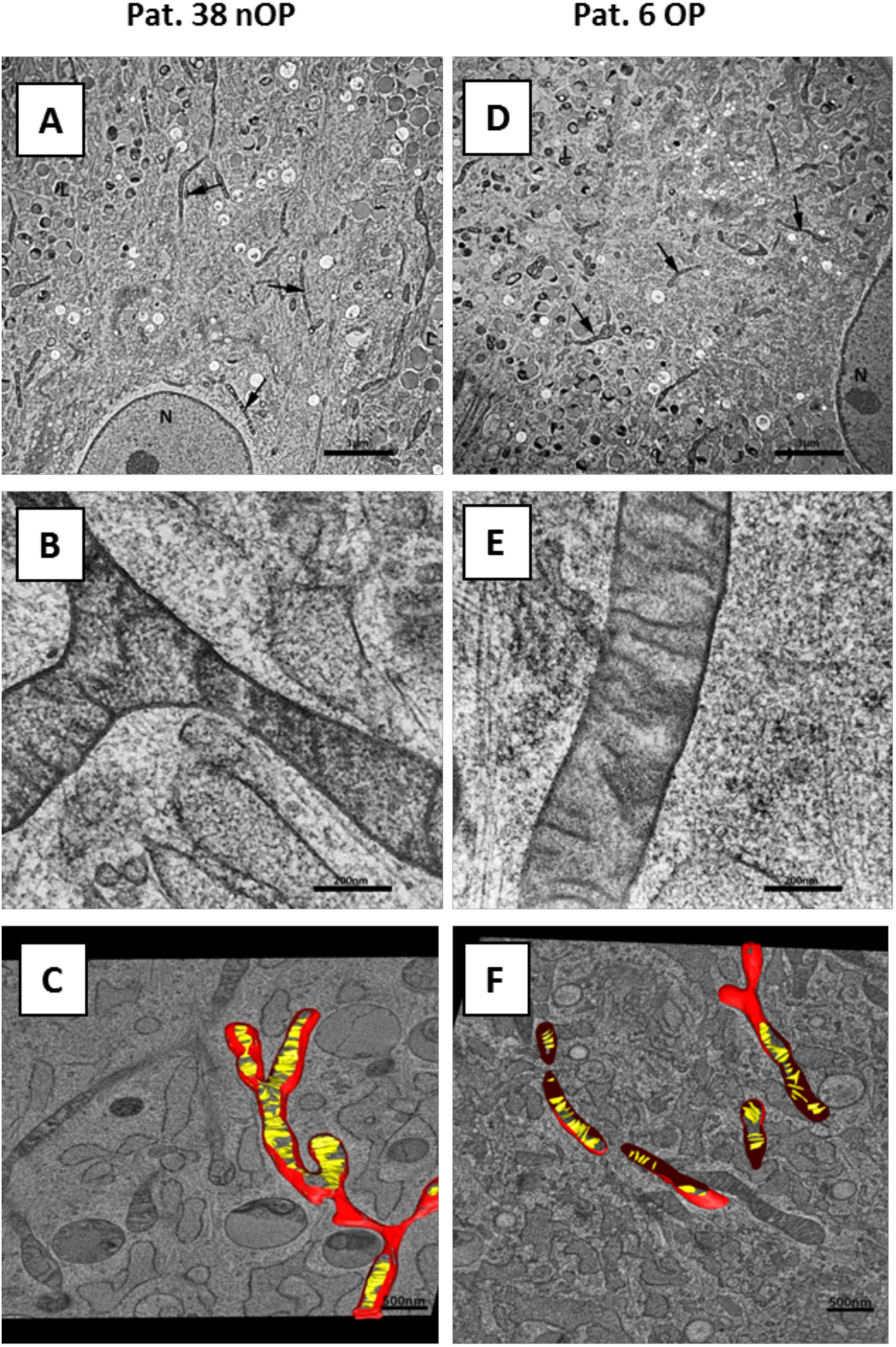
High resolution electron micrograms of MSC harvested from non osteoporotic (A, B, C) and osteoporotic (D, E, F) donors. Figs. A and D show an overview of organelles in the cytoplasm and the predominant perinuclear localization of mitochondria indicated by arrows. N denotes nucleus and L (with arrowhead) Lyso-autophagosomes. Figs. B and E display highly magnified mitochondria with inconspicuous cristae and a well preserved outer double membrane. Figs. C and F show 3D reconstructions based on an electron tomogram of a single, branched, threadlike mitochondrion with cristae shown in yellow and mitochondrial membrane in red.

### 3.3. Respiratory assessment of cellular metabolism

The coupling control protocol applied to intact hBMSCs indicated that there is neither a significant difference detectable in the extent of ROUTINE respiratory activity between cells derived from healthy subjects and osteoporotic donors, nor was ROUTINE respiration any different in the osteosarcoma cell line MG-63 (Fig. 3A). Similarly, there was no significant difference between the three groups in their LEAK respiration, although there was a 1.6-fold higher rate in the OP and MG-63 cells as compared to nOP cells. In contrast, the maximum respiration observed at optimum uncoupler concentration applied to intact cells was somewhat higher in nOP than in OP and MG-63 cells, but given the relatively high variability there was also no statistical significance of these differences.

**Fig. 3:**
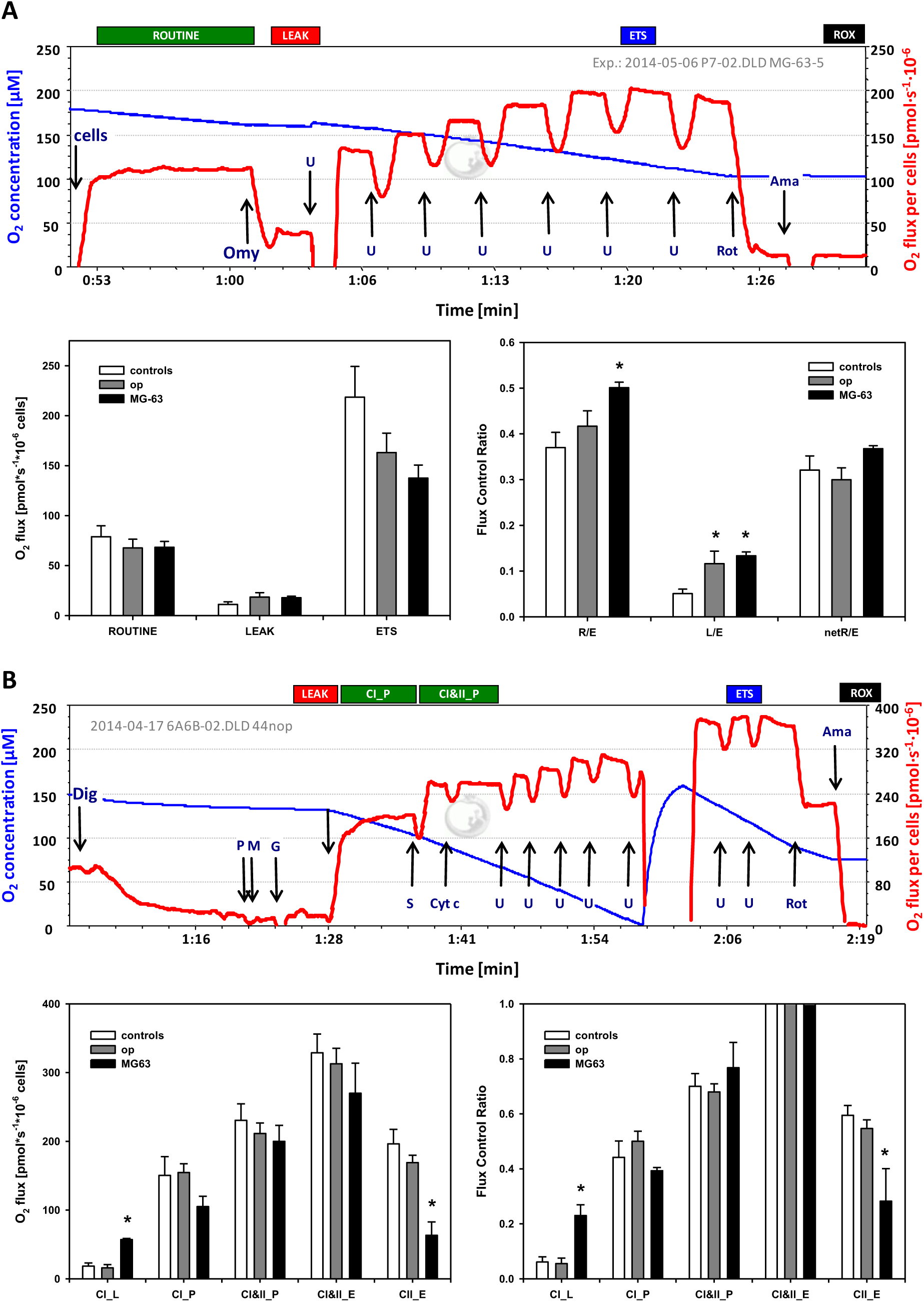
Respiratory characteristics of hBMSCs from non-osteoporotic and osteoporotic donors and of the osteosarcoma cell line MG-63. A: respiratory evaluation of intact cells in a coupling control protocol. Upper image: representative trace of a respiratory experiment with intact MG-63 cells. Oxygen concentration is depicted as a blue line (left y-axes [μM]) and derived oxygen flux per million cells is shown in red (right y-axis [pmols·^-1^ml·^-1^]). Bars above the traces denote respiration in ROUTINE state, in the LEAK state, in the uncoupled state in which electron transfer system capacity, ETS, is attained, and residual oxygen consumption, ROX. Lower panel: summarized results of cell number specific respiratory rates presented as means ± SE of 6, 7, and 9 independent cultures, each measured in duplicate (left) and of flux control ratios calculated from these data showing the ratios of ROUTINE respiration to ETS capacity (R/E), of LEAK respiration to ETS capacity (L/E), and of net ROUTINE respiration to ETS capacity ((R-L)/E)) (right). B: evaluation of respiratory characteristics of permeabilized hBMSCs and MG-63 cells applying a substrate-uncoupler-inhibitor titration (SUIT) protocol. Upper image: representative trace of a respiratory experiment with hBMSCs. Image shows oxygen concentration (blue line, left y-axes [μM]) and derived oxygen flux per million cells (red line, right y-axis [pmols·^-1^ml·^-1^]) and additions of digitonin (Dig), of respiratory substrates pyruvate (P), malate (M) and glutamate (G), of ADP (D), succinate (S), of cytochrome c (Cyt c), of uncoupler CCCP up to maximum respiratory activity (U) and of CI inhibitor rotenone (Rot) and of CIII inhibitor antimycin A (Ama). Bars above the traces denote respiration in LEAK state, CI-linked substrate supported OXPHOS (CI_P), combined CI and CII-linked substrates supported OXPHOS (CI&II_P), electron transfer system capacity (ETS), and residual oxygen consumption (ROX). Lower panel: summarized results of cell number specific respiratory rates presented as means ± SE of 6, 7, and 4 independent cultures, each measured in duplicate (left) and of flux control ratios calculated from these data by normalizing rates to ETS supported by combined CI‐ and CII-linked substrates (CI&II_E) (right).

Besides the assessment of absolute respiratory rates per cell number, it may also be useful to compare various flux control ratios derived from these rates, as this enables normalization of fluxes independently of variations in e.g. cell size. When using ETS capacity as a reference state and expressing ROUTINE and LEAK respiration as a fraction of ETS, it can be seen that ROUTINE respiration operates at approximately 40% of ETS capacity in hBMSCs of healthy and osteoporotic donors, but at about 50% in the ostosarcoma cell line. The ratio between LEAK respiration and ETS capacity, L/E, shows that in nOP cells LEAK accounts for only 6% of ETS capacity and to about 12% in OP cells and MG-63. The resulting flux control ratio for netROUTINE, (ROUTINE-LEAK)/ETS, an indicator of the fraction of respiration coupled to ATP production relative to ETS capacity, indicates that there is no difference among groups in this regard.

Evaluation of respiration in permeabilized cells indicated that there were also no differences detectable between OP and nOP cells in the various substrate and coupling states examined (Fig. 3B). Thus, rates were almost identical in LEAK respiration, in respiration coupled to ATP production with primarily complex-I linked substrates (CI_P) or with substrates supporting convergent complex-I and ‐II supported OXPHOS (CI&II_P), as well as in convergent complex-I and ‐II supported uncoupled respiration (CI&II_E) and uncoupled complex-II linked activity (CII_E). In comparison, MG-63 cells displayed significantly higher LEAK respiration and lower uncoupled complex-II linked respiration. Overall, OP and nOP cells appear energetically indistinguishable, whereas the osteosarcomic cell line shows some significant differences.

Given the uneven gender distribution of donors of nOP and OP cells we evaluated whether the absence of respiratory differences between both groups might be related to this. In both intact and permeabilized cells, the one female nOP sample showed somewhat higher respiration than most male nOP cells, while the single male OP sample was slightly below average of the female dominated OP group. Based on only one sample in each group this does not allow to rule out a gender bias, but as these differences were diminishing rather than enlarging any trend towards a group difference, they render a general gender bias rather unlikely.

## 4. Discussion

The exact etiology of osteoporosis is far from being understood and may be related to functional disturbances of fully differentiated cells involved in bone metabolism and preservation of structure, or in the altered capability of bone-marrow stem cells to adequately support the renewal of differentiated cells and thereby maintain a balance between bone formation and resorption [14]. Based on findings supporting the latter idea [4,15], we investigated whether altered energetics of bone-marrow derived stem cells could underlie expression of the osteoporotic phenotype, investigating mitochondrial structure and function.

We first confirmed that the cells isolated from bone marrow were actually pluripotent stem cells by establishing some standard characteristics. We demonstrate expression patterns of surface markers typical of mesenchymal stem cells and the absence of markers characteristic for hematopoietic cells. We show that the cells adhere to a plastic surface, and we confirmed the potential to differentiate, upon adequate stimulation, along the adipogenic, chondrogenic and osteogenic lineages. All these criteria were met by cells from both OP and nOP patients, indicating that general characteristics of hBMSC are preserved in OP cells.

Next, we investigated cellular ultrastructural features using electron microscopy technique supporting optimum preservation of organelle structure. We found that mitochondrial appearance was unaltered in OP cells and did not show any of the typical signs of mitochondrial dysfunction such as swelling or enlargement of the cristae. Further, there was also no obvious change in mitochondrial number, although we admittedly did not quantify this in any strict manner. In contrast, changes in mitochondrial size and distribution have been reported for brain and peripheral cells of patients with bipolar disorders [16], in the number [17] and appearance of mitochondria of cancer cells [18], and in the cristae morphology and size of mitochondria of neurons from patients with Alzheimer’s disease [19,20]. Noteworthy, we saw a high number of lyso-autophagosomes in the hBMSCs indicative of dynamic membrane turnover, but this was seen in OP and nOP-derived cells and is thus clearly not related to osteoporosis.

Our respirometric analysis of intact and permeabilized hBMSC indicated that no major differences exist between cells from nOP and OP patients in terms of ROUTINE respiration, LEAK respiration, ETS capacity, or the relative contribution of CI and CII to OXPHOS or maximum respiratory activity. Some differences observed for LEAK and ETS in intact cells were not significant and upon permeabilization of the cell membrane, i.e. under conditions where substrate supply to the respiratory system was not a rate limiting parameter, OP cells appeared completely indistinguishable from nOP cells (Fig. 3B). This differs from findings by Oliveira et al., who reported that bone marrow cells from obese mice had impaired respiratory capacity, reflected as both reduced ROUTINE respiration of intact cells and as reduced OXPHOS capacity in permeabilized cells [21]. Similarly, lymphocytes from patients with Alzheimer’s disease showed significantly diminished OXPHOS capacity [22].

A comparison of our OP and nOP cells with an established osteosarcomic cell line used in the present study and with various other cell types investigated with identical methods and summarized by Pesta and Gnaiger suggests that respiratory characteristics of hBMSC fall well into the range determined for other primary cells and cell lines of both human and murine origin [13]. Thus, despite the notion that stem cells residing e.g. in the bone marrow or the uterus rely heavily on anaerobic glycolysis, reflecting their hypoxic environment, the present data suggest that this does not become apparent as an obvious peculiarity of their respiratory characteristics [9,23]. In line, as summarized by Lonergan et al.[9], both lower [24] as well as higher rates of oxygen consumption [25] were reported for undifferentiated stem cells as compared to their differentiated counterparts. It should be noted, however, that oxygen-dependent characteristics may only become apparent at physiological oxygen levels, as also pointed out by Lonergan [9].

In summary, our data suggest that despite the dynamic changes of cell metabolism described to be associated with differentiation processes of stem cells, there is no evidence that mitochondrial alterations manifested as structural and/or energetic alterations contribute to the development of osteoporosis, at least not the form of the disease typically expressed in the elderly patients that were the subjects of the present study. It should be mentioned that more subtle changes, such as altered signaling or changes in substrate affinities affecting mitochondrial energetics as observed e.g. in tissues from Diabetes patients have not been examined here and the existence of this type of alterations cannot be ruled out on the basis of our present data [26].

## Acknowledgements

We thank M.W. Hess for helpful discussions and providing the electron microscopic infrastructure. We also acknowledge P.J. Richards for sharing the MG-63 cell line as well as fruitful discussions on stem cells. Furthermore, we thank S. Sopper for providing the infrastructure of the FACS core facility.

